# Germplasm Collections and Morphological Studies of *Andropogon gayanus-Andropogon tectorum* Complex in Southwestern Nigeria

**DOI:** 10.1101/2022.09.10.507395

**Authors:** Funmilola Mabel Ojo, Chinyere Constance Nwokeocha, Julius Olaoye Faluyi

## Abstract

Morphological studies were carried out on collections of *Andropogon gayanus-Andropogon tectorum* complex from Southwestern Nigeria. This is with the view to providing full characterization of the accession of the two species of *Andropogon* and elucidating their population dynamics.

Morphological data from selected accessions of *A. gayanus* and *A. tectorum* from different parts of Southwestern Nigeria were collected and characterized using an adaptation of the *Descriptors for Wild and Cultivated Rice (Oryza* spp), Biodiversity International. Preliminary morphological description of the accessions were carried out at the point of collection. Garden populations were raised from the vegetative parts of some accessions and maintained in Botanical Garden of the Obafemi Awolowo University, Ile-Ife. The data obtained were subjected to inferential tests and Duncan’s multiple range test.

The results revealed a distinct distribution pattern of the two species of *Andropogon* in the area of study which suggests a south-ward migration of *Andropogon gayanus*, that is from the northern vegetational zones of Nigeria to the southern ecological zones. The migration of *A. gayanus* around Igbeti with occasional occurrence of *A. tectorum* along the roadsides without any distinct phenotypic hybrid, and Budo-Ode in Oyo State is established as the southern limit of the spread of *A. gayanus*. This migration of *A. gayanus* to the South is not an invasion but a slow process. There was no *A. gayanus* encountered in Osun, Ondo, Ekiti and Ogun States. *Andropogon gayanus* and *Andropogon tectorum* do not only emerge from the rootstocks rapidly but can also produce independent propagules by rooting at some nodes. The plants can spread by means of these propagules even if it does not produce sexual or apomictic seeds. This potential for vegetative propagation in addition to the perennial habit confer considerable advantage for colonization by the *Andropogon gayanus-Andropogon tectorum* complex.

## INTRODUCTION

The genus *Andropogon* Linn. is a fairly large genus of the grass family, Poaceae, belonging to the tribe Andropogoneae (Olorode 1984; Hutchinson & Dalziel 1972). *Andropogon* is a pantropical genus of grasses of about 29 species almost confined to the tropical and warm temperate regions of the world, frequently forming an important part of the savanna vegetation in the tropics. *Andropogon* is represented by about 14 species in Nigeria (Lowe, 1989) although Stanfield (1970) had reported about 12 species. The genus is composed of annual and perennial species frequently with tall culms, and leaf blades which can be linear to lanceolate or ovate. The spikelets occur in pairs at each node of the raceme, each pair consisting of a pedicellate and a sessile spikelet. The sessile spikelet is bisexual, the pedicellate is unisexual male (Hutchinson and Dalziel, 1972). They are articulated in such a way that at maturity, the spikelets, pedicel and internodes all break apart leaving no central inflorescence stalk. The sessile spikelet bears a prominent awn which is flexed at an angle to the vertical axis of the glumes. A distinct colour difference exists between the two arms of the awn (Clayton, 1969; Stanfield, 1970).

*Andropogon gayanus* Kunth is a tall, tufted perennial grass that grows taller than 3 m. It has various tillers and abundant foliage especially during the rainy season (Chlleda and Crowder, 1982). It forms a significant part of the vegetation of many savanna areas throughout Africa south of the Sahara, including South Africa. It is a polymorphic species. In Nigeria, four main varieties were recognized (Clayton, 1962). These are: var. *gayanus* (var. *genuinus*) Hack; var. *bisquamulatus* (Hochst) Hack var. *squamulatus* (Hochst) Stapf and var. *tridentatus*. Bowden (1963) considered var. *tridentatus* as split from var. *bisquamulatus* thus recognizing only three varieties. *A. gayanus* is widespread and abundant in the Northern and Southern Guinea Savanna, as well as, in the drier areas of the derived savanna whereas, *A. tectorum* occupies vast areas in the derived savanna, preferring moderate shade (Stanfield, 1970). However, certain areas in the derived savanna support the growth of both species equally well (Okoli and Olorode, 1983). *Andropogon gayanus* is propagated by seeds, which are broadcasted or planted in rows and vegetatively by splitting the tufts. It is relatively free of major pests and diseases and is resistant to grazing and burning. These make it a useful grass for supporting a large number of ruminant animals in Northern Nigeria. It is also one of the high-yielding grasses in West Africa (Bogdan, 1977; Pagot, 1993). The economic importance of *Andropogon gayanus* for livestock grazing is that it is very palatable when young and serves as basic materials for woven houses. *Andropogon gayanus* is a highly-productive grass, which increases fuel loads, produces intense, late dry season fires which seriously damage native woody species; it is also useful as forage in permanent pastures grazed by ruminants. The stems are used for thatching and, when flattened, for weaving coarse grass mats as well as sometimes being planted for erosion control and soil restoration.

*Andropogon tectorum* Schum. & Thonn. on the other hand, is a perennial grass; caespitose. Culms can be 200 – 300 cm long without nodal roots or with prop roots. Ligules are eciliate membrane or a ciliolate membrane, 1–2 mm long. Leaf blade base tapers to the midrib and bears false petiole. Leaf-blades are lanceolate; 30 – 45 cm long; 10 – 25 mm wide; flaccid; light-green, apex is acuminate.

*Andropogon gayanus* and *A. tectorum* are important natural fodder grasses in Nigeria. They are also useful in crop rotation, thatching and mat-making (Bowden, 1963) and offer an interesting opportunity for ecological, cytogenetic and evolutionary studies.

The major components of this study are: germplasm collection and morphological studies. There is paucity of information about the taxonomic and cytogenetic studies of this complex species, besides the first attempts by Okoli (1978). The dynamics of the population are not fully known. Besides, for about two decades and more, a re-assessment of these two agriculturally-important grasses is overdue. As already stated, observations from pre-survey revealed that the localization of *A. gayanus* to the North may no longer be true. There were larger population of plants looking very much like *A. gayanus* which have invaded the derived savanna ecosystem in Nigeria. Plants that look intermediate also occur abundantly in locations in the derived savanna ecosystem. It therefore becomes necessary to address the problems above through germplasm collection and characterization of the two species involved in this complex.

## MATERIALS AND METHIODS

Field trips for plant collections covered agro-ecological zones of the following states: Osun, Ondo, Ogun, Oyo and Ekiti as shown in Figure 1. Whole plants of *Andropogon gayanus* and *Andropogon tectorum* were collected from different locations in Southwest, Nigeria. Accession numbers were given to the specimens. Seeds were also collected for planting and preservation. Garden populations were raised from the vegetative parts of some accessions and they were also maintained in the Botanical Garden of the Obafemi Awolowo University, Ile-Ife, Osun State. The accessions were nurtured to maturity and used for this study. Table 1 shows the locations, coordinates, descriptions of site and collectors of the accessions.

**Table 1:** Accessions of *Andropogon gayanus* (Kunth) - *Andropogon tectorum* (Schum & Thonns) Studied and their Sources.

| ACCN | LOCATIONS | COORDINATES | DESCRIPTIONS OF SITE | COLLECTORS |
| --- | --- | --- | --- | --- |
| AG 1 | Asawo, Oyo State | N 08° 59.848` E 04° 17.836` | Ruderal location, regular regrowth forest | Ojo, Faluyi, Azeez and Abraham |
| AG 2 | Budo-Ode, Oyo State | N 08° 20.420` E 04° 14.039` | Derived Savanna | '' |
| AG 3 | Between Ogbomosho and Oko, Oyo State | N 08° 59.854` E 04° 17.835` | Ruderal location, regular regrowth forest | '' |
| AG 4 | Igbeti 1, Oyo State | N 08° 24.722` E 04° 15.467` | Dry water course, populated by heavy mat of grasses and sedges; chimeric lawn | '' |
| AG 5 | Igbeti 2, Oyo State | N 08° 24.723` E 04° 15.467` | Dry water course, populated by heavy mat of grasses and sedges; chimeric lawn | '' |
| AG 6 | Ogbomosho, Oyo State | N 08° 34.794` E 04° 14.669` | Regular regrowth forest with the presence of multiple grass species | '' |
| AT 1 | OAUTHC Road, Osun State | N 07° 30.870` E 04° 33.065` | Ruderal location with dwarf morphotypes propagating from crevices | Ojo, Faluyi and Nwokeocha |
| AT 2 | O.A.U. Religious Centre, Osun State | N 07° 30.703` E 04° 33.065` | Ruderal on lateritic soil in company of other grasses, Asteraceae | '' |
| AT 3 | O.A.U. International School, Osun State | N 07° 31.793` E 04° 33.261` | Expanse of lateritic soil with close communities of *Andropogon tectorum* | '' |
| AT 4 | Ife-Ibadan road, Osun State | N 07° 22.774` E 04° 01.497` | At the fringe of road divider, surrounded by other grasses | Ojo and Faluyi |
| AT 5 | Ondo road, Along Ore, Ondo State | N 07° 49.468` E 04° 52.444` | Ruderal on lateritic soil | Ojo |
| AT 6 | Aladura, Ogun State | N 07° 30.068` E 04° 27.885` | Ruderal on lateritic soil | Ojo and Faluyi |
| AT 7 | Omu-Ayede road, Ekiti State | N 07° 54.374` E 05° 20.167` | Ruderal location with a cluster of *Panicum maximum* and stands of *Chromolana odorata* | Ojo, Faluyi, Matthew and Abraham |
| AT 8 | Itaji-Oye Road, Ekiti State | N 07° 50.833` E 05° 20.661` | Ruderal location under a tree of *Parkia biglobosa* and *Alchornea laxiflora* | Ojo, Faluyi, Matthew and Abraham |
| AT 9 | Ayede-Oye Road, Ekiti State | N 07° 50.438` E 05° 20.733` | Solitary plant, leaves are broad and usually short; in mesic location | '' |
| AT 10 | Erinmo Road, Ekiti State | N 07° 38.203` E 04° 51.755` | Ruderal location on lateritic soil in company of other grasses, Asteraceae | '' |

ACCN – Accession, AG- *Andropogon gayanus*, AT- *Andropogon tectorum*

**Fig. 1:**
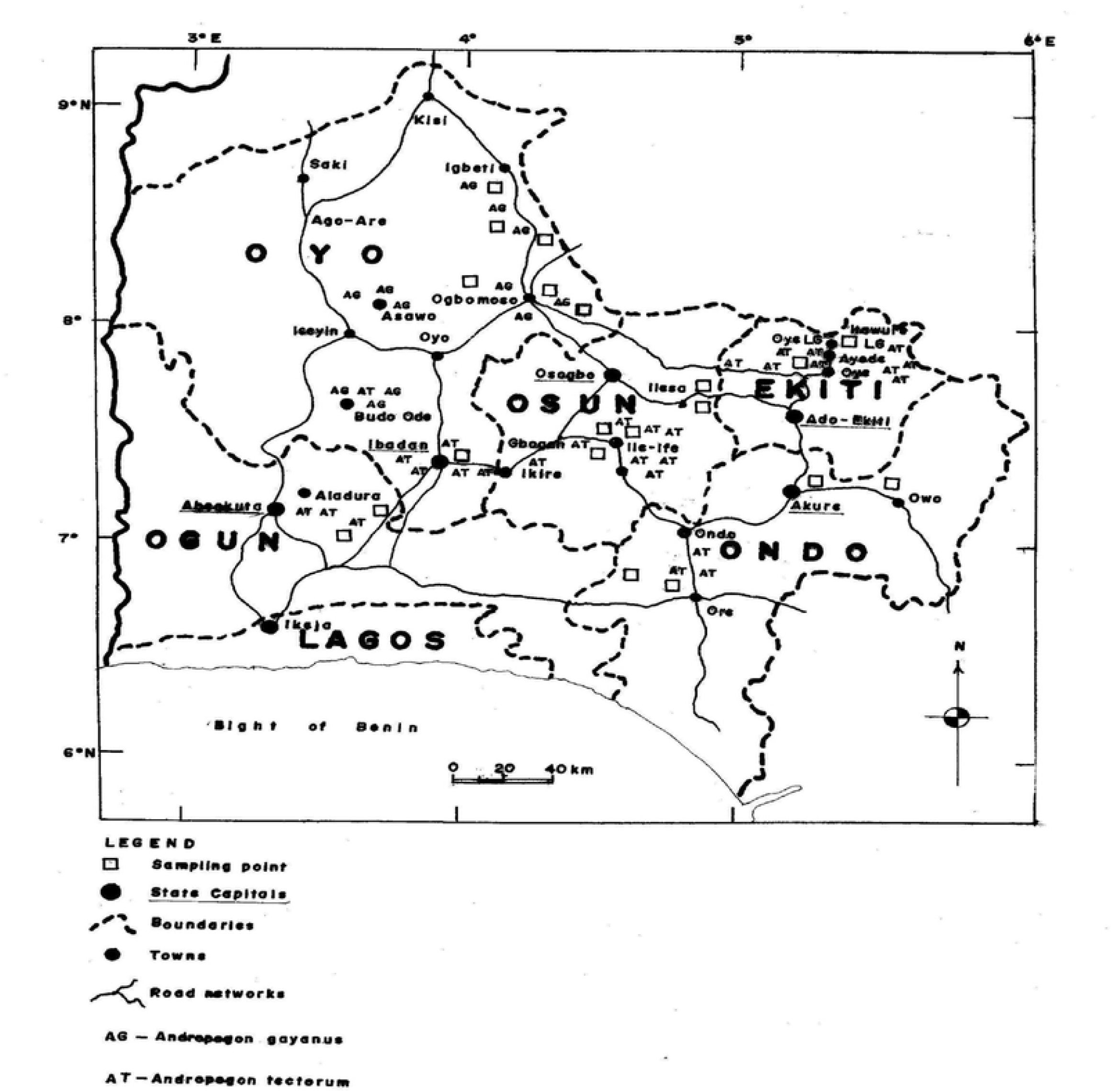
Map Showing Collection Sites.

Quantitative, as well as, qualitative morphological characters were observed and measured for each species as required. The leaf area was estimated. Some quantitative data were measured using the millimeter graph paper method. The characters studied were described using an adaptation of the *Descriptors for Wild and Cultivated Rice (Oryza spp)*, Biodiversity International (2007) as follow:

1. Culm characteristics – number, height (measured on the main culm from soil level to the base of the panicle after heading) and pigmentation.
2. Habit – annual or perennial
3. Leaf characters – this includes the length and breadth of the flag leaf and leaf below flag, flag leaf angle, pubescence, keel characteristics (absence, presence and prominence), and length and hairiness of petiole.
4. Inflorescence characters – (floral morphology: stigma colour, awn colour, length of upper glume of pedicellate floret, length of upper glume of sessile floret, type of ovary etc.), number of raceme pairs that is possible on a flowering axis, length of raceme, counts of spikelet on a raceme pair, length and breadth of spikelets
5. Seed – seed shape (oval, oblong, round, elongated), seed length, caryopsis colour.

## RESULTS

### Spatial Distribution of the Two Species Studied

The population density of *Andropogon gayanus* in Igbeti is high. The occurrence is in large clusters with solitary stands of *Andropogon tectorum* at the peripheries and occasional small clusters in open spaces. The roadside pattern is similar with clusters of *A. gayanus* interspersed by lone stands of *A. tectorum*. That is the typical formation of the two species along the road down-south except that the relative proportion of the plants switched in favour of *A. tectorum*. Budo-Ode (N 08º 20.420’ E 04º 14.039’) is the southern limit beyond which *A. gayanus* was a rare occurrence.

The two plant species were presented as pure morphological samples throughout the region surveyed. The variation in hairiness observed in *A. tectorum* samples is probably an environmental effect which can be associated with closeness to the road and dry environments as observed on the Ife-Ibadan road, the Obafemi Awolowo University Campus and several locations in Ekiti State.

### Morphological Studies

#### *Andropogon gayanus* Plate 1, Tables 2, 3, 4 and 5

##### Habitat/ Habit

The accessions of *Andropogon gayanus* were found on dry water courses populated by a heavy mat of grasses and sedges, chimeric lawn, on the road fringes. They thrive well on all lands where there is enough soil for them to grow. They grow well in the open. Some are very large in population and spreading mainly through root stocks and prop roots from the nodes (Plate 1). They compete well with other plants.

**Table 3:** Qualitative Floret Attributes of the Accessions of *Andropogon gayanus-Andropogon tectorum* Studied.

| ACCN | SPIKELET COLOUR | SPIKELET SHATTERING | NATURE OF END OF SPIKELET | STIGMA COLOUR | ANTHER COLOUR | AWN COLOUR |
| --- | --- | --- | --- | --- | --- | --- |
| AG1 | Brown | + | Blunt | Creamy white | Yellowish-brown | Light green |
| AG2 | " | + | " | Light purple | " | " |
| AG3 | " | + | " | " | " | " |
| AG4 | " | + | " | Creamy white | " | " |
| AG5 | " | + | " | " | " | " |
| AG6 | " | + | " | Light purple | " | " |
| AT1 | " | + | " | " | " | Light purple |
| AT2 | " | + | " | Creamy white | " | " |
| AT3 | " | + | " | " | " | " |
| AT4 | " | + | " | " | " | " |
| AT5 | " | + | " | " | " | " |
| AT6 | " | + | " | " | " | " |
| AT7 | " | + | " | " | " | " |
| AT8 | " | + | " | Light purple | " | " |
| AT9 | " | + | " | Creamy white | " | Light green |
| AT10 | " | + | " | " | " | " |

**ACCN** – Accession, **AG-** *Andropogon gayanus*, **AT-** *Andropogon tectorum*.

**Table 4:** Quantitative of Some Vegetative Features of the Accessions of *Andropogon gayanus-Andropogon tectorum* Studied.

| AC. | CULM |  |  |  | LEAF CHARACTERS |  |  |  |  |
| --- | --- | --- | --- | --- | --- | --- | --- | --- | --- |
|  | No of Culm | Length (cm) | Density (cm) | Leaf area (cm<sup>2</sup>) | Flag leaf Length (cm) | Flag leaf Breadth (cm) | Leaf below flag leaf Length (cm) | Leaf below flag leaf Breadth (cm) | Petiole Length (cm) |
| AG1 | 71 | 209.41±2.25 | 1.24±0.07 | 342.45±18.70 | 44.52±1.20 | 6.14±0.16 | 52.58±0.93 | 5.16±0.09 | 23.00±0.89 |
| AG2 | 75 | 198.97±2.98 | 1.43±0.04 | 328.54±20.87 | 45.1±1.22 | 5.28±0.18 | 51.48±1.10 | 5.44±0.13 | 21.8±1.02 |
| AG3 | 72 | 206.81±2.86 | 1.66±0.06 | 358.89±19.45 | 44.82±1.59 | 5.88±0.19 | 52.82±1.3 | 5.14±0.16 | 19.20±1.02 |
| AG4 | 79 | 200.28±1.18 | 1.25±0.02 | 342.75±13.05 | 44.84±1.62 | 5.96±0.20 | 51.90±0.60 | 5.88±0.12 | 21.80±1.11 |
| AG5 | 74 | 203.47±2.44 | 1.14±0.07 | 372.89±17.55 | 46.08±1.68 | 5.84±0.21 | 51.00±1.01 | 5.92±0.07 | 19.20±0.86 |
| AG6 | 76 | 193.37±2.45 | 1.43±0.03 | 403.55±20.22 | 47.14±1.07 | 5.72±0.22 | 53.5±1.37 | 5.00±0.30 | 22.60±1.72 |
| AT1 | 69 | 114.47±3.70 | 1.36±0.06 | 497.45±16.55 | 40.82±0.60 | 5.86±0.14 | 54.14±1.96 | 4.86±0.14 | 50.40±3.66 |
| AT2 | 75 | 191.68±3.80 | 1.62±0.05 | 422.65±18.14 | 52.32±1.00 | 5.72±0.26 | 56.28±0.28 | 5.00±0.30 | 33.20±3.98 |
| AT3 | 77 | 192.69±3.90 | 1.57±0.07 | 451.45±20.55 | 51.58±1.00 | 5.67±0.26 | 55.28±0.28 | 4.98±0.30 | 33.50±3.98 |
| AT4 | 79 | 198.67±3.90 | 1.53±0.07 | 444.67±12.78 | 46.93±2.10 | 5.63±0.40 | 56.5±0.68 | 5.65±0.23 | 33.50±2.99 |
| AT5 | 71 | 189.77±3.00 | 1.48±0.05 | 414.76±15.19 | 46.42±1.30 | 5.28±0.22 | 57.86±0.77 | 4.96±0.09 | 32.80±3.97 |
| AT6 | 74 | 199.61±3.70 | 1.58±0.04 | 470.78±16.55 | 49.00±0.90 | 5.28±0.25 | 56.66±0.55 | 5.04±0.12 | 29.60±2.98 |
| AT7 | 76 | 242.79±14.50 | 1.63±0.06 | 405.78±18.19 | 44.76±1.10 | 5.92±0.21 | 51.72±1.30 | 5.28±0.19 | 23.30±1.39 |
| AT8 | 78 | 220.56±9.60 | 1.55±0.06 | 418.76±17.12 | 45.98±1.00 | 5.92±0.31 | 50.60±0.75 | 5.18±0.28 | 19.80±0.73 |
| AT9 | 76 | 236.42±11.85 | 1.78±0.07 | 407.78±20.11 | 48.20±1.20 | 5.94±0.17 | 53.98±1.14 | 5.30±0.15 | 22.40±1.72 |
| AT10 | 99 | 309.50±6.90 | 2.95±0.06 | 345.76±20.12 | 56.04±1.18 | 6.86±0.23 | 57.86±0.77 | 6.06±0.50 | 36.80±2.85 |

**AC** – Accession, **AG**- *Andropogon gayanus*, **AT**- *Andropogon tectorum*.

**Table 5:** Quantitative Reproductive Features of the Accessions of *Andropogon gayanus-Andropogon tectorum* Studied.

| **ACCN** | **MNRP** | **MLRP**<br>**(cm)** | **MNSP** | **MLSP**<br>**(cm)** | **MBSP**<br>**(cm)** | **LUGS**<br>**(cm)** | **LUGP**<br>**(cm)** | **SL (cm)** |
| --- | --- | --- | --- | --- | --- | --- | --- | --- |
| AG1 | 8.90±0.12 | 17.76±0.11 | 38.00±0.90 | 7.80±0.12 | 2.20±0.09 | 3.40±0.19 | 3.40±0.19 | 2.70±0.12 |
| AG2 | 8.20±0.24 | 18.14±0.27 | 38.60±0.75 | 7.20±0.25 | 2.25±0.11 | 3.30±0.20 | 3.50±0.16 | 2.50±0.16 |
| AG3 | 8.60±0.19 | 18.50±0.23 | 37.60±0.98 | 7.30±0.15 | 2.30±0.09 | 3.40±0.19 | 3.50±0.22 | 2.70±0.11 |
| AG4 | 8.20±0.12 | 18.66±0.21 | 31.60±0.55 | 7.40±0.19 | 2.30±0.12 | 3.50±0.22 | 3.50±0.16 | 2.30±0.12 |
| AG5 | 9.30±0.13 | 18.48±0.19 | 37.60±0.60 | 7.30±0.30 | 2.10±0.10 | 3.40±0.19 | 3.60±0.19 | 2.70±0.20 |
| AG6 | 9.50±0.22 | 18.28±0.24 | 39.50±0.90 | 7.40±0.29 | 2.20±0.12 | 3.60±0.18 | 3.40±0.17 | 2.60±0.19 |
| AT1 | 7.50±0.19 | 11.20±0.25 | 23.30±0.44 | 5.19±0.19 | 1.20±0.12 | 2.50±0.16 | 2.40±0.16 | 1.95±0.05 |
| AT2 | 9.90±0.24 | 13.26±0.36 | 24.20±0.30 | 5.60±0.19 | 1.40±0.06 | 2.70±0.12 | 2.60±0.19 | 1.90±0.06 |
| AT3 | 9.85±0.24 | 13.18±0.36 | 23.80±0.34 | 5.58±0.19 | 1.42±0.06 | 2.68±0.13 | 2.55±0.19 | 1.87±0.06 |
| AT4 | 8.90±0.33 | 11.20±0.25 | 24.00±0.22 | 5.80±0.12 | 1.50±0.08 | 2.60±0.19 | 2.30±0.12 | 1.85±0.06 |
| AT5 | 8.50±0.20 | 12.40±0.38 | 29.75±0.61 | 5.40±0.24 | 1.43±0.07 | 2.75±0.14 | 2.90±0.13 | 1.94±0.00 |
| AT6 | 9.60±0.15 | 12.84±0.38 | 29.60±0.25 | 5.80±0.12 | 1.55±0.06 | 2.80±0.12 | 2.80±0.12 | 1.90±0.06 |
| AT7 | 9.50±0.18 | 15.82±0.27 | 29.90±0.75 | 6.20±0.12 | 2.00±0.08 | 3.20±0.12 | 3.20±0.12 | 2.20±0.12 |
| AT8 | 10.60±0.18 | 15.70±0.46 | 30.00±0.63 | 6.20±0.12 | 2.10±0.06 | 3.30±0.11 | 3.20±0.11 | 2.20±0.12 |
| AT9 | 11.70±0.18 | 15.60±0.27 | 30.40±0.75 | 6.30±0.20 | 2.10±0.13 | 3.10±0.10 | 3.20±0.12 | 2.10±0.17 |
| AT10 | 14.30±0.12 | 22.86±0.40 | 37.20±0.49 | 7.25±0.10 | 2.40±0.06 | 3.85±0.18 | 3.60±0.17 | 2.35±0.10 |

**ACCN** – Accession, **AG** – *Andropogon gayanus*, **AT** - *Andropogon tectorum*, **MNRP**- Mean Number of raceme pairs, **MLRP**- Mean length of raceme pairs, **MNSP**- Mean number of spikelets, **MLSP**- Mean length of spikelets, **MBSP**- Mean breadth of spikelets, **LUGS**- Length of upper glume sessile, **LUGP**- Length of upper glume pedicellate, **SL** – Seed length.

**Table 6:** Principal Components Analysis for Quantitative Morphological Characters of the Accessions of *Andropogon tectorum-Andropogon gayanus* Studied.

| Variables | Component 1 | Component 2 | Component 3 |
| --- | --- | --- | --- |
| CH | 0.16 | 0.50 | 0.26 |
| LFL | -0.05 | 0.60 | 0.06 |
| BFL | 0.17 | -0.11 | 0.83 |
| PL | -0.16 | 0.43 | -0.15 |
| LLBFL | 0.21 | 0.34 | 0.01 |
| BLBFL | -0.26 | 0.11 | 0.19 |
| MNRP | 0.31 | 0.09 | -0.13 |
| MLRP | 0.33 | 0.02 | -0.03 |
| MNSP | 0.31 | 0.03 | -0.32 |
| MLSP | 0.32 | -0.05 | -0.13 |
| MBSP | 0.30 | -0.16 | 0.19 |
| LUGS | 0.33 | 0.09 | 0.04 |
| LUGP | 0.33 | 0.09 | -0.01 |
| SL | 0.32 | -0.14 | -0.10 |
| **Eigen Values** | **8.69** | **2.15** | **0.92** |
| **Variance** | **6.54** | **1.23** | **0.18** |
| **Proportion** | **0.62** | **0.15** | **0.07** |

**ACCNS**- Accessions, **CH**- Culm height, **LFL**- Length of flag leaf, **BFL**- Breadth of flag leaf, **LLBFL**- Length of leaf below flag leaf, **BLBFL**- Breadth of leaf below flag leaf, **PL**- Petiole length, **MNRP**- Mean number of raceme pairs, **MLRP**- Mean length of raceme pairs, **MNSP**- Mean number of spikelets, **MLSP**- Mean length of spikelets, **MBSP**- Mean breadth of spikelets, **LUGS**- Length of upper glume sessile, **LUGP**- Length of upper glume pedicellate, **SL**- Seed length.

The accessions studied were all perennials. New tillers regenerated from root stocks of old plants at the beginning of the rains. Tussocks recover well after fire with rapid development of new tillers.

##### Plant Types

The accessions studied were mainly erect and open, some tillers lodging to the ground at maturity, most especially after heavy wind. Plant growth is about 193 – 209 cm high, the culm is relatively fat (1.52 – 1.92 cm) covered with hairy leaf sheath; no pigmentation. The plants are robust and low to heavy-tillering (23 - 51 tillers per plant). There is expression of hairiness in the leaf sheath, petiole and lamina. Leaf texture is foamy, colour of the outer leaf sheath is green, no pigmentation, hairy. The leaf is pubescent on both sides, linear-lanceolate, acute; leaf can be up to 40 cm long and about 2.2 cm in breadth hairy at base. Flag leaf angle is acute (45º). Ligule is an eciliate membrane; presence of keel but not prominent. The petiole can grow up to 28 cm, hairy. The stem is pubescent and green. The seed is oblong with length between 2 cm and 3 cm, Caryopsis colour is dark-brown. The spikelets shatter easily, this habit is aided by strong wind; the spikelets ends are blunt. The mean number of spikelets ranged from 31.60 - 39.50; there were just slight variations in the measurements of the spikelets. The stigma colour is either creamy-white or light-purple; awn colour is light-green and the anther is yellowish-brown.

#### *Andropogon tectorum* Plate 2, Tables 2, 3, 4 and 5

##### Habitat/Habit

The accessions of *Andropogon tectorum* were found on road dividers, mesic locations, ruderal on lateritic soil in company of other grasses in close proximity to *Panicum maximum*. They thrive well on all lands where there is enough soil for them to grow. They grow well in the open. Some are very large in population and are propagated mainly through root stocks and propagules generated by rooting through the nodes (Plate 2). They compete well with other plants and survive.

The accessions studied were all perennials. Observations in the field showed that regeneration and spreading is mainly through rootstocks; fire does not destroy the rootstocks. Tussocks recover well after fire with rapid development of new tillers.

##### Plant Types

The accessions studied were mainly erect and open, new tillers regenerated from root stocks of old plants immediately after the first rain. Plant grows to about 114 - 330 cm tall, the culm is relatively fat (1.65 – 1.98 cm) in diameter, hairless, purple pigmented, with whitish powdery substances on the culm. The plants are robust and heavy-tillering (31 - 69 tillers per plant). AT1, collected along OAUTHC (N 07º 30.870’ E 04º 33.065), possesses stout culms with profuse nodal roots whereas the plant is short (miniature). Leaf pubescence is intermediate; some leaves are glabrous while some are hairy but not as dense as in *A. gayanus*. Leaves are alternate, lanceolate with acuminate apices; leaf can be up to 50 cm in length and about 2.8 cm in breadth, glabrous at base, outer leaf sheath is purple, inner leaf sheath is green, Flag leaf angle is acute (45º), ligule is membranous; a keel is present and prominent; the petiole can be glabrous or pubescent and long (can attain a height of 62 cm). Stems are glabrous, green and purple; young stems are covered by white substances which could be removed by touch (it runs off on being touched) with the exception of few ones which when young has hairy leaf blade**s**.

The exposed part of the stem has purple pigmentation. However, the accessions collected for *A. tectorum* fell into two field forms: tall, heavy-tillering with broad leaves and big culm having whitish powdery substances, purple pigmentation and the stout, low-tillering with narrow leaves and thin culm, no pigmentation. These forms occurred most times.

The seed is oblong with length between 1.75 cm and 2.5 cm, caryopsis colour is light-brown. Spikelets shatter easily aided by strong wind; the ends of the spikelets are blunt. The mean number of spikelets ranged from 23.30 to 37.30; spikelets are longest (7.25 cm) and broadest (2.40 cm) in AT10, Kiwani, a collection from the Itawure-Erinmo Road (N 07 º38.203’ E 04 º51.755’) in Ekiti State. They are shortest (5.18 cm) and narrowest (1.20 cm) in AT1 collected along OAUTHC road (N 07º 30.870’ E 04º 33.065), Ile-Ife, Osun State. Generally, the stigma colour is creamy-white or light-purple; awn colour is light-purple and anther is yellowish-brown.

Principal Components Analysis (Figure 2) shows that a collection of plants in the *Andropogon gayanus-Andropogon tectorum* complex can easily be separated into two, based on proximity using the quantitative morphological characters employed in the analysis. The accessions collected along OAUTHC road and Kiwani collected on the Itawure-Erinmo Road are far from each other and far from the other groups. The accessions from Aladura and Ife-Ibadan Road are closely related to the accessions from OAU Religious Centre and OAU International School. Omu-Ayede and Ayede-Oye Road accessions are closely related and more related to the accessions from Itaji-Oye Road. All the accessions of *A. gayanus* from Asawo, Budo-Ode, Igbeti; between Ogbomosho and Oko, Ogbomosho, aggregate closely to each other which showed close relationship among them.

**Fig. 2:**
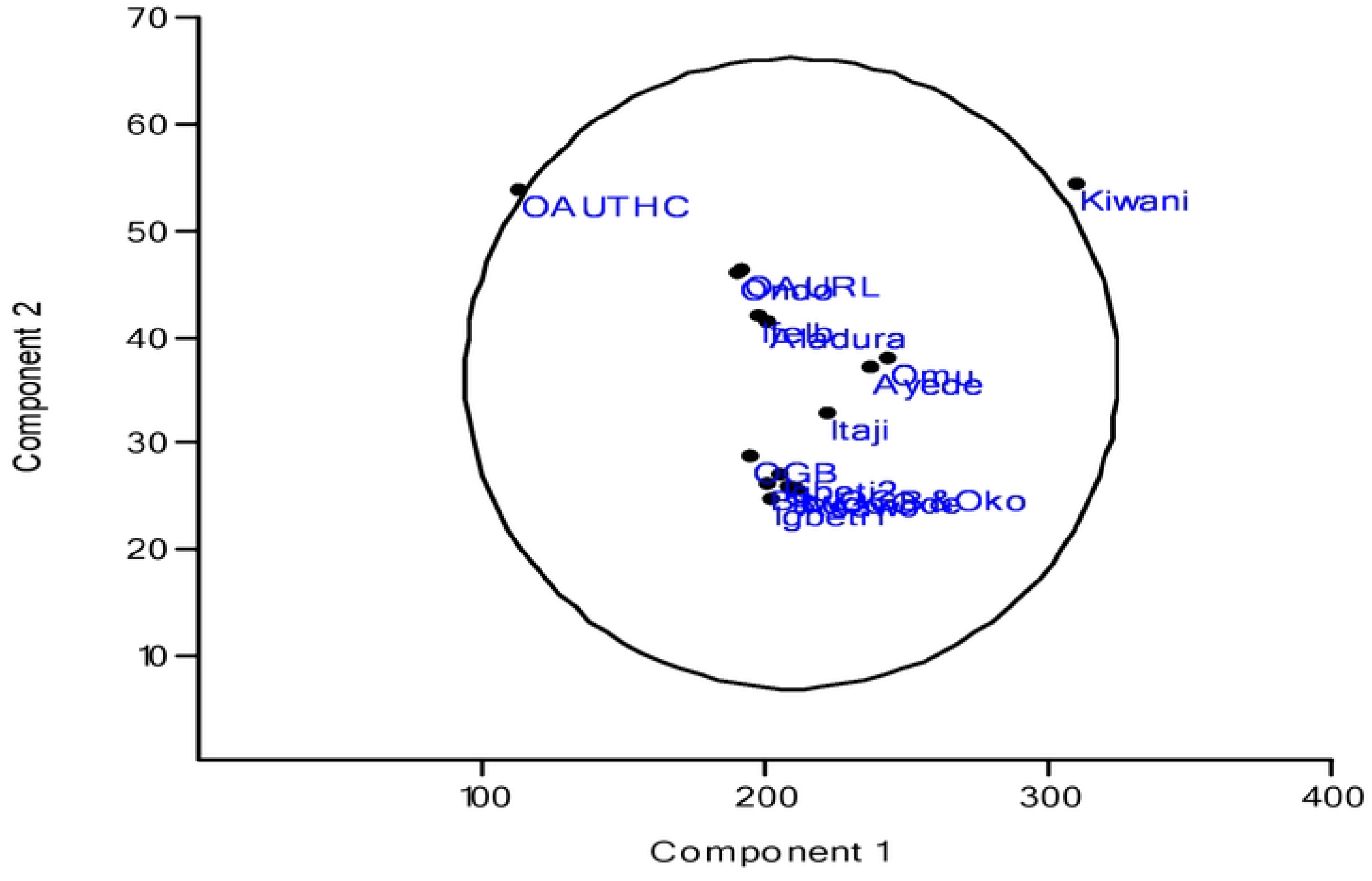
Scattered Diagram of the Accessions Studied Based on Quantitative Morphological Characters.

## DISCUSSION

This survey has revealed the distribution pattern of the two species of *Andropogon* in the area of study which conforms to a south-ward migration of *Andropogon gayanus* from the northern vegetational zones of Nigeria to the southern ecological zones. Okoli (1978) made collections of *A. gayanus* from Niger State, Kwara State, Enugu State and the outskirts of Ogbomosho on the Ilorin road. This survey is forty two years after that of Okoli (1978) and Budo-Ode in Oyo State has been established as the southern limit of the spread of *A. gayanus*. The above observation suggests that the migration of *A. gayanus* to the South is not an invasion but a slow process. The trajectory is ruderal, the establishment exploits open spaces and the process is facilitated by anthropogenic factors: movement of people, vehicles, cattle-rearing, farming activities, bush-burning, etc. It is reasonable to assume that the migration of *A. gayanus* to the South is due to a similar ecological preferences to *A. tectorum* suggesting that they may have a distinct genomic identity. *Andropogon gayanus* has established well in Igbeti but the presence of *A. tectorum* is clearly evident. The slow rate of spread is also consistent with seed dispersal as a means of transfer. The study on seed germination shows that *A. gayanus* is between 43% and 60% and the seed set is good despite spikelet shattering which is a serious economic problem for seed production in agreement with Sanada *et al*., (1998).

The study ensured exploration of a large area of the southwest and extensive collections from a wide range of habitats, characterization and elucidation of the species complex. The results obtained in this study have helped to shed more light on the population dynamics of *Andropogon gayanus-Andropogon tectorum* complex in southwestern Nigeria.

The field survey on *A. tectorum* revealed two field forms: tall, heavy tillering with broad leaves and big culm (heavy bloom) having whitish powdery substances, purple pigmentation and the stout, low tillering with narrow leaves and thin culm (moderate or sparse bloom) no pigmentation.; The result of the meiotic study (though not reported in this paper) and the population dynamics in the population of *A. gayanus* and *A. tectorum* have largely been understood.

The distribution of *Andropogon gayanus* and *Andropogon tectorum* are largely distributed in Southwestern Nigeria. They are found as ruderals, on waste lands, abandoned and cultivated farm lands, regular regrowth forest, expanse of lateritic soil, water courses, grazing grounds populated by heavy mat of grasses and sedges, chimeric lawns on the road fringes and road dividers where there is sufficient soil to aid their growth. The habits of the two species are mainly perennials, they are colonizers which have succeeded in displacing other plant species in their major areas of occurrence. These observations correspond with the findings of several authors like Burkill (1935); Lowe (1989); Akobundu and Agyakwa (1998).

In most floras, exomorphology has been used to delimit taxa. Morphological attributes constitute the orthodox tool of the taxonomist because of their diagnostic value in taxonomic evaluation as opined by Ayodele and Olorode, (2005). The morphology of an organism is not simply as a result of accident but represents the result of a long evolution of successive adaptations in living organisms to their environment. Morphological study is known to play a crucial role in species identification and documentation. The Qualitative morphological results of this study largely correspond with those of Hutchinson and Dalziel (1972) and Lowe (1989). Morphological characters have also been used by other researchers such as Smith and Ashton, (2006); Adedeji and Illoh, (2005); Adedeji and Faluyi, (2006) to enhance the taxonomy of different taxa Saheed and Illoh (2011) successfully used morphological characters to separate *Senna* and *Chamae-crista* from *Cassia* despite some overlaps (indication of closeness of the genera and species) recorded in some of the characters.

*Andropogon gayanus* and *Andropogon tectorum* not only emerge from the rootstocks rapidly but can also produce independent propagules by rooting at some nodes. The plants can spread by means of these propagules even if it does not produce sexual or apomictic seeds. This potential for vegetative propagation, in addition to the perennial habit, confer considerable advantage for colonization by the *Andropogon gayanus-Andropogon tectorum* complex. *Andropogon gayanus* was not as successful on the field compared to *A. tectorum*. It was observed that lodging took place on the accessions collected from Igbeti when they were raised on the field in the Botanical Garden of the Obafemi Awolowo University, Ile-Ife probably because of the weather condition of the new environment for example, high amount of rainfall compared to diminished rainfall of Igbeti. Biomass and plant architecture at the point of collection could also account for this. Adegbite (1991) reported a similar strategy in *Aspilia africana* (Pers) C. D. Adams at maturity which was primarily based on its ability to extend through vegetative propagation. Few plant stand could spread, mostly by vegetating through an advanced enlarging rootstock, especially during the growing season.

In the *A. gayanus–A. tectorum* complex, interspecific and intraspecific vegetative features vary and overlap to a large extent. The taxonomic delimitation of the components of this complex into species had been largely based on some vegetative features such as leaf width, pigmentation, hairiness, petiole form and length, flowering scape. Certain inflorescence characteristics like hairiness of spikelets have also been used in the taxonomy of these species especially at various level in *A. gayanus*. Again, considerable variation has been observed. Plants in this complex appear to vary more within each species than between species. In all accessions studied, morphological characters adopted separate Kiwani, from others because Kiwani featured large biomass typified by a very high number of tillers, broad but short leaves, and dense expression of hairiness.

The collections of *A. gayanus* showed that the hairiness of certain organs is independent of other parts. Some plants possess glabrous or nearly-glabrous leaf sheaths but have densely hairy lamina and petioles; some possess hairy leaf sheaths and petioles. These variations in the frequency and degree of hairs on various plant organs were observed.

The key issues involved in the distribution pattern of the species and population dynamics within and among the two species of *Andropogon* studied can be summarized as follows:

1. the dominant species around Igbeti is *Andropogon gayanus* with occasional occurrence of *Andropogon tectorum* along the roadsides without any distinct phenotypic hybrid. This trend continued as far South as Budo-Ode (N 08º20.420’ E 04º 14.039’) in Oyo State.
2. there is southward migration of *A. gayanus* around Igbeti with occasional occurrence of *A. tectorum* along the roadsides without any distinct phenotypic hybrid, and Budo-Ode in Oyo State has been established as the southern limit of the spread of *A. gayanus*. No *A. gayanus* was encountered in Osun, Ondo, Ekiti and Ogun States.
3. propagation of *Andropogon gayanus* and *Andropogon tectorum* are mainly through root stocks and aerial roots from the nodes

## CONCLUSION

The major elements of the population dynamics of the *Andropogon gayanus-Andropogon tectorum* complex have been elucidated in this study. Seed propagation will remain the major process of the spread of this complex, hybridization will play a crucial role as the ratio of the two species improve as a result of the establishment of *Andropogon gayanus* in its southward expansion.

## ACKNOWLEDGEMENT

The research was financially supported by Bill Dahl Graduate Student Research Grant Award (2019), Botanical Society of America, United States of America and Academic Staff Union of Universities (ASUU) Research Grant Award in support of Doctoral thesis for 2019/2020 Academic Session.

---

**Plate 1:**
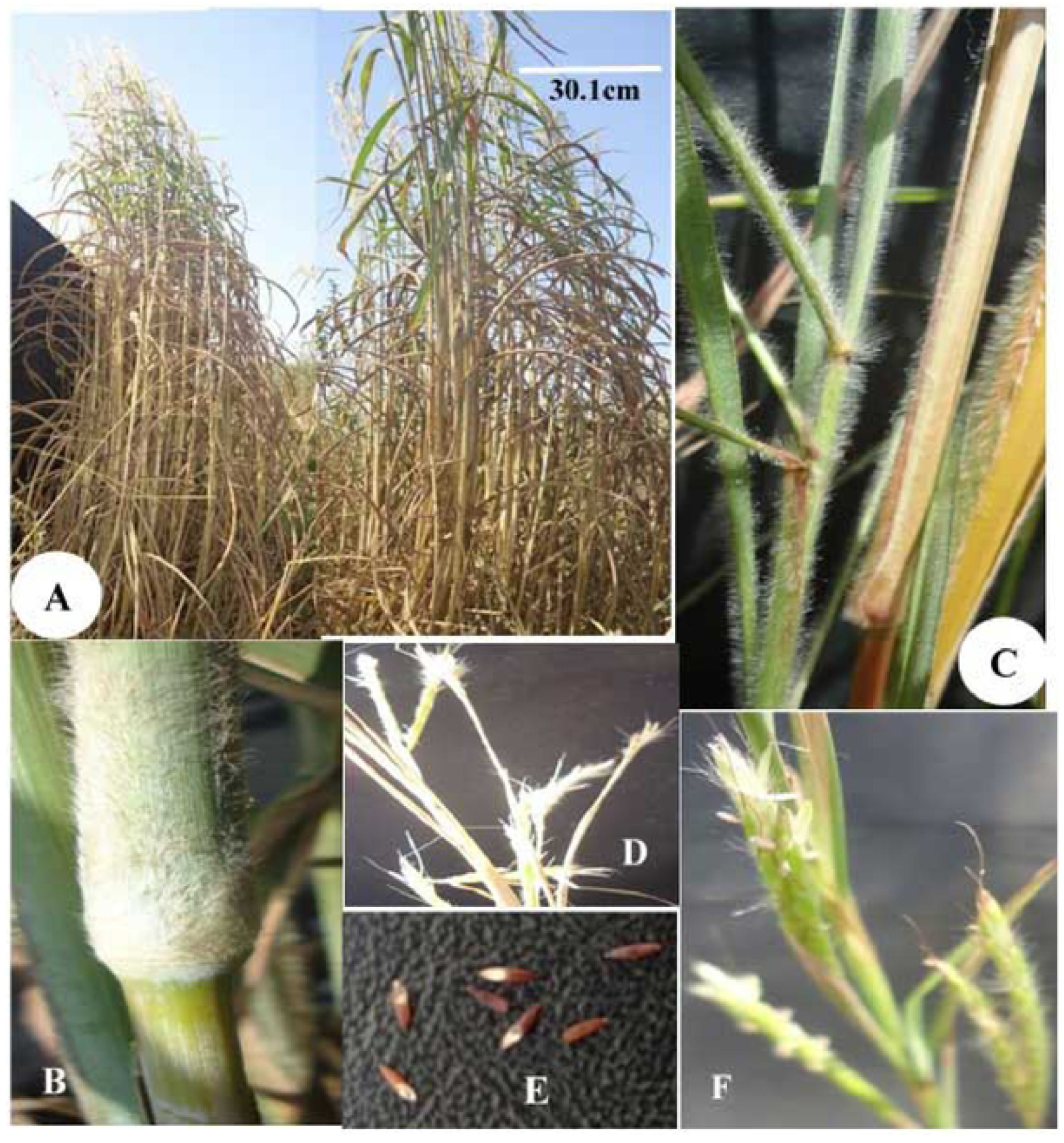
Morphological features of *Andropogon gayanus*. A- Plant form and adaptation B- internode C- Hairy leaf sheath D- Raceme pairs E- Seeds F- Spikelet

**Plate 2:**
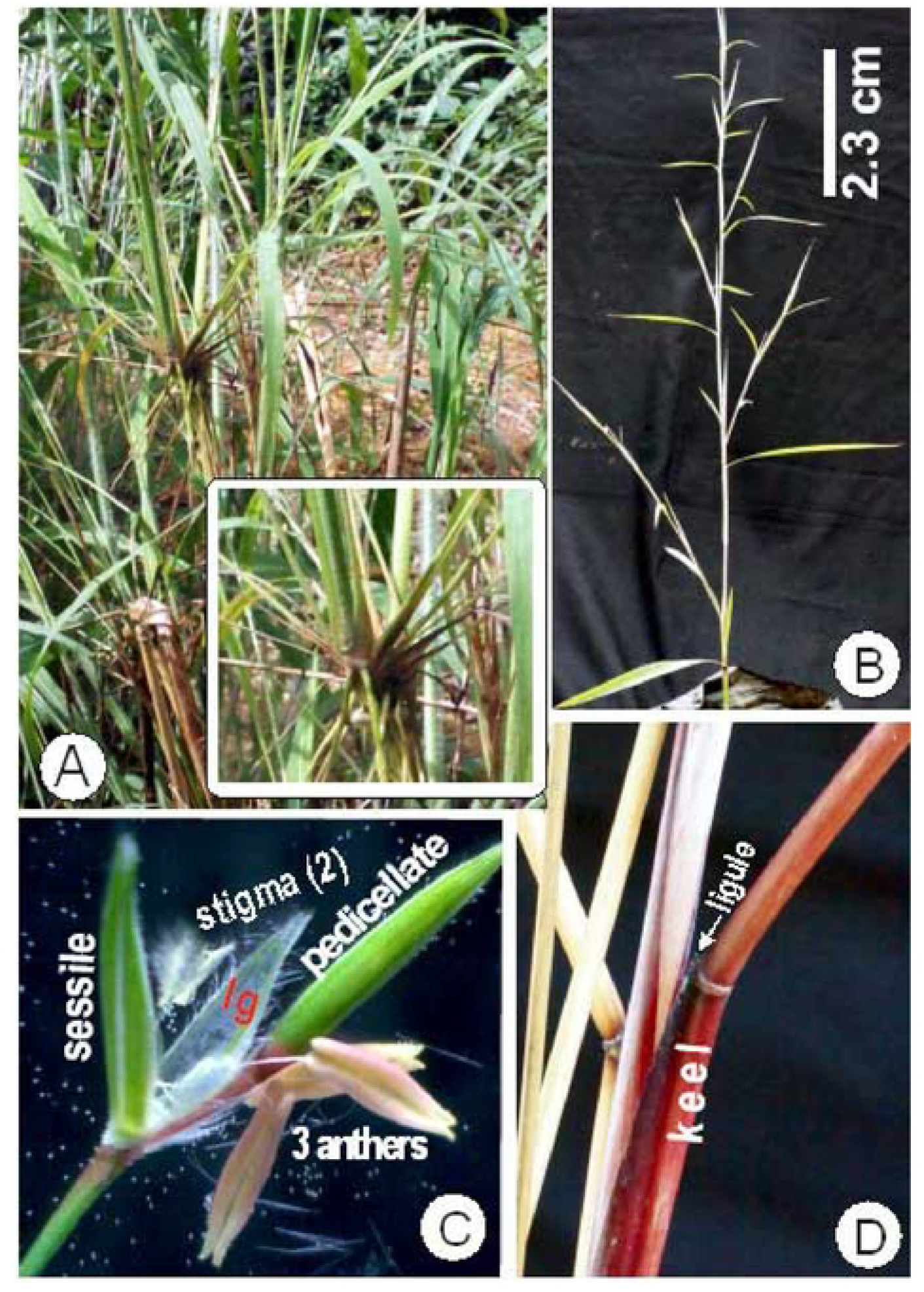
Morphological Features of *Andropogon tectorum*. A- Plant form and adaptation (Insect-Rooting at the node) B- Flowering Scape C- Spikelet D- The leaf showing sheath, keel and ligule ; lg – lower glume

